# Dynamic cerebellar network organization across the human menstrual cycle

**DOI:** 10.1101/2020.05.29.123869

**Authors:** Morgan Fitzgerald, Laura Pritschet, Tyler Santander, Scott T. Grafton, Emily G. Jacobs

## Abstract

The cerebellum contains the vast majority of neurons in the brain and houses distinct functional networks that constitute at least two homotopic maps of the cerebrum. While the functional organization of the human cerebellum has been characterized, the influence of sex steroid hormones on intrinsic cerebellar network dynamics has yet to be established. Here, we investigated the extent to which endogenous fluctuations in estradiol and progesterone alter functional cerebellar networks at rest in a woman densely sampled over a complete menstrual cycle (30 consecutive days). Edgewise regression analysis revealed negative associations between sex hormones and cerebellar coherence, with progesterone showing more pronounced negative associations relative to estradiol. Graph theory metrics probed sex hormones’ influence on topological brain states, revealing relationships between sex hormones and intra- and inter-network integration in Ventral Attention, Dorsal Attention, and Somato-Motor Networks. Together, these results suggest that the intrinsic dynamics of the cerebellum are intimately tied to day-by-day changes in sex hormones.

---

Although its Latin name means “little brain”, the human cerebellum contains nearly four times as many neurons as the cerebral cortex, with the posterior and lateral regions greatly expanded in humans relative to apes (Andersen et al., 1992). Engagement of the cerebellum during cognitive control tasks challenge the classic notion that the cerebellum is solely involved in motor coordination; rather, it appears to coordinate a broad range of higher-order cognitive functions (Marek & Dosenbach, 2018; Ramnani et al., 2006). Multiple closed-loop circuits between the cerebellum and cortex, including non-motor regions of the prefrontal cortex (PFC; Middleton & Strick, 1994; Kelly & Strick, 2003), provide an anatomical basis for cerebellar involvement in cognition, including learning, memory, and decision making (Timman et al., 2010; Mandolesi et al., 2001; Harrington et al., 2004). Thus, the tradition of branding the cerebellum as a purely motor-associated region is becoming increasingly obsolete.

Allen and colleagues (2005) demonstrated the utility of using functional magnetic resonance imaging (fMRI) to assess functional synchrony between the cerebellum and the cerebral cortex, finding that low-frequency signal fluctuations in the cerebellum correlate with signal fluctuations in subcortical, parietal, and frontal regions. Topographically distinct fronto-cerebellar circuits involving the dorsolateral PFC, medial PFC, and anterior PFC have since been identified (Krienen & Buckner, 2009). A seminal fMRI study by Buckner and colleagues (2011) revealed that the cerebellum houses at least two complete homotopic maps of cortical networks. The cerebellum displays representations of major functional brain networks including the Default Mode Network (DMN), Frontal Control Network (FCN), Somato-Motor Network (SMN), Dorsal Attention Network (DAN), Ventral Attention Network (VAN), and Limbic Network (Buckner et al., 2011).

Accumulating evidence implicates the cerebellum as a major site of sex steroid hormone action. In rodents, the cerebellum exhibits *de novo* synthesis of estradiol and progesterone during critical developmental periods (Tsutsui et al., 2011; Sakamoto et al., 2001) and demonstrates a rich expression of estrogen receptors (ER) and progesterone receptors (PR) in adulthood (Shughrue et al., 1997; Price & Handa, 2000; Guerra-Ariaza et al., 2002; Pang et al., 2013), suggesting that sex hormones not only influence the formation of cerebellar neuronal circuitry but also modulate cerebellar functioning later in life. The vast majority of Purkinje cells, the major output units of the cerebellum, densely express ERβ (Price & Handa, 2000; Shughrue et al., 1997), and estradiol has been shown to improve cerebellar memory formation by enhancing long-term potentiation and augmenting cerebellar synapse formation (Andreescu et al., 2007). Although much attention has been paid to sex hormones’ ability to regulate spinogenesis, synaptic plasticity, and neural activity in cortex (Hara et al., 2015; Woolley & McEwen, 1993; Frick et al., 2018), sex hormones’ role in the cerebellum is now gaining increased recognition.

Despite preclinical evidence that sex hormones regulate cerebellar function, human studies are lacking. Across a typical human menstrual cycle, spanning 25-30 days, women experience a ~12-fold increase in estrogen and an ~800-fold increase in progesterone. These *in vivo* changes in sex hormones have been shown to modulate brain structure, task-evoked cortical activity, and performance on cognitive tasks (Berman et al., 1997; Jacobs & D’Esposito, 2011; Lisofsky et al., 2015; Galea et al., 2017). However, most menstrual cycle studies sparsely sample women at discrete timepoints (e.g. 2 days), obscuring the rhythmic changes in hormone production across a complete cycle. The field of network neuroscience has begun to use dense-sampling methods to probe the dynamic properties of the human brain over unprecedented timescales (Poldrack et al., 2015; Barth et al., 2016; Pritschet et al., 2019). In a recent dense-sampling study from our group, a woman underwent 30 consecutive days of brain imaging and venipuncture across a complete menstrual cycle, revealing estradiol and progesterone’s ability to modulate widespread patterns of connectivity across the cortex (Pritschet et al., 2019). Given the sensitivity of the cerebral cortex to endogenous fluctuations in sex steroid hormones (Weis et al., 2019; Pritschet et al., 2019; Taylor et al., 2020; Jacobs & Goldstein, 2018) and accumulating evidence for sex hormone action in the cerebellum (Brinton et al., 2008; Hedges et al., 2012), here we tested the hypothesis that sex hormones impact the intrinsic dynamics of cerebellar circuits.

In this dense-sampling, deep-phenotyping study, we examined whether day-by-day variation in sex hormones across a complete menstrual cycle modulates cerebellar functional connectivity and cerebellar network topologies. Results reveal that estradiol and progesterone are strongly associated with daily variation in coherence across the cerebellum and both intra- and inter-network integration, providing insight into how sex hormones shape the intrinsic dynamics of the human cerebellum.

## Results

A healthy, naturally-cycling female (author L.P.; age 23) underwent venipuncture and MRI scanning for 30 consecutive days. The full dataset consists of daily mood, diet, and behavioral assessments; task-based and resting-state fMRI; structural MRI; and serum assessments of pituitary gonadotropins and ovarian sex hormones (Pritschet et al., 2019). Neuroimaging data, daily behavioral assessments, and analysis code will be accessible upon publication.

### Endocrine assessments

Analysis of daily sex hormone (by liquid-chromatography mass-spectrometry) and gonadotropin (by chemiluminescent immunoassay) concentrations confirmed the expected rhythmic changes of a typical menstrual cycle. All hormones fell within normal ranges (**Table S1**), with a total cycle length of 27 days. Serum levels of estradiol and progesterone were lowest during menses (day 1-4) and peaked in the late follicular (estradiol) and late luteal (progesterone) phases (**Figure 1**). Progesterone concentrations surpassed 5ng/mL in the luteal phase, signaling an ovulatory cycle (Leiva et al., 2015).

**Figure 1.**
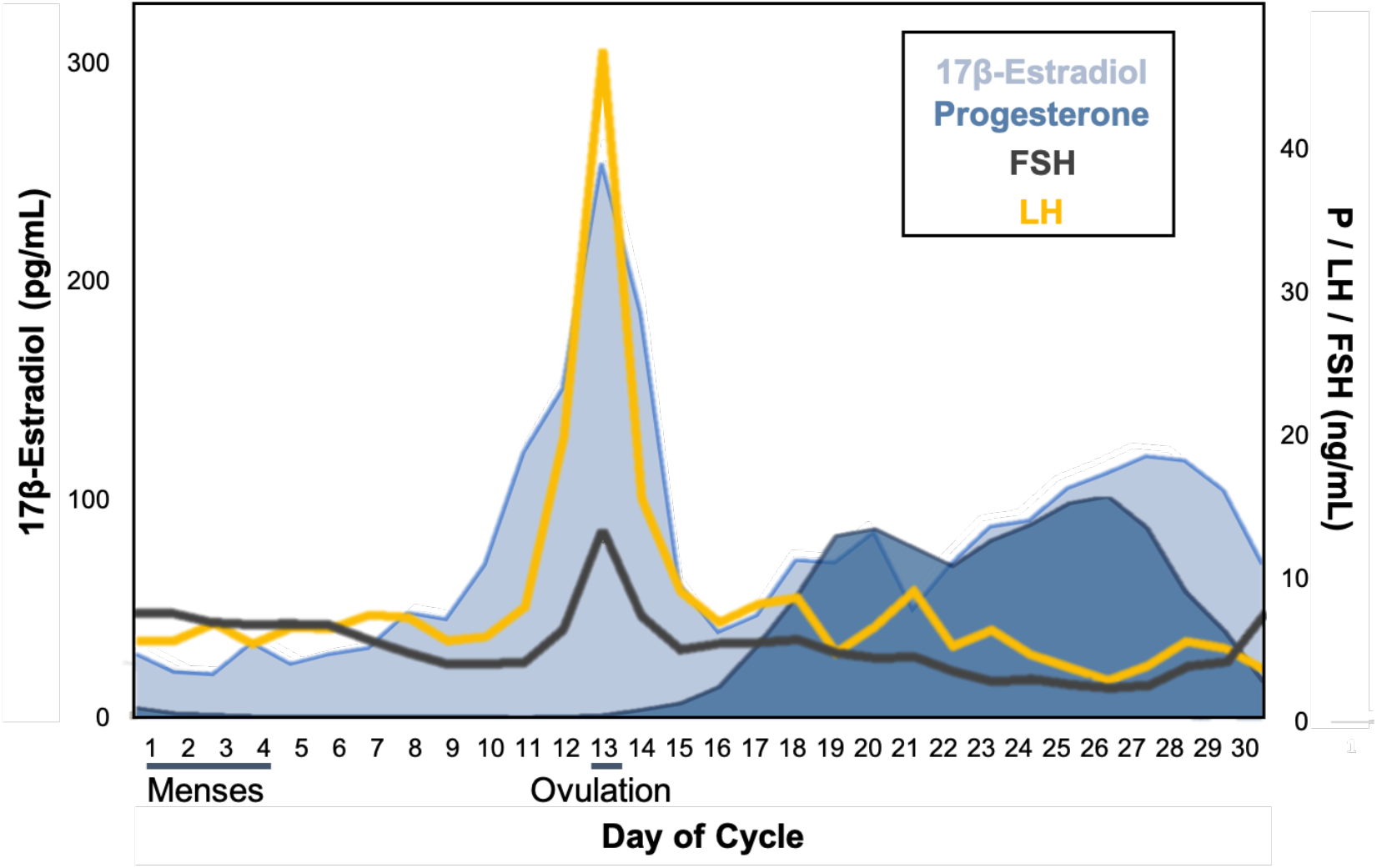
Participant’s hormone concentrations plotted by day of cycle. 17β-estradiol, progesterone, luteinizing hormone (LH), and follicle stimulating hormone (FSH) concentrations fell within standard ranges. Adapted from Pritschet et al. 2019.

### Temporal dependencies between sex hormones and edgewise connectivity

To begin, we tested the hypothesis that cerebellar functional connectivity at rest is associated with intrinsic fluctuations in estradiol and progesterone in a day-by-day fashion. Given the pronounced expression of PR within the cerebellum and the ability of progesterone to augment inhibitory responses within cerebellar neurons (Pang et al., 2013, Smith et al., 1987; Brinton et al., 2008), we predicted decreases in cerebellar functional connectivity as progesterone concentrations increase across the cycle, in keeping with results previously found in the cerebrum (Pritschet et al., 2019). Further, we predicted estradiol would augment cerebellar coherence (Pritschet et al., 2019). For each session, the cerebellum was parcellated into 99 nodes from the Ren atlas that were then mapped to the Buckner seven-network atlas (Ren et al., 2019; Buckner et al., 2011). A summary time-course was extracted from each node, data were temporally filtered, and 99 × 99 functional association matrices were derived via magnitude-squared coherence (FDR-thresholded at *q* < .05; see **Methods and Materials** for full description of preprocessing and connectivity estimation). Next, we specified edgewise regression models, regressing coherence against estradiol and progesterone over the 30-day study. Data were *Z*-scored prior to analysis and models were thresholded against empirical null distributions generated through 10,000 iterations of nonparametric permutation testing. Results reported below survived a conservative threshold of *p* < .001.

In keeping with our predictions, progesterone yielded a widespread pattern of robust inverse associations across the cerebellum, such that whole-cerebellar coherence decreased as progesterone concentrations rose (**Figure 2A**). Next, the average magnitude of brain-hormone association was summarized by network (using the Buckner seven-network parcellation; **Figure 2C**), revealing that all networks demonstrate some degree of positive associations over time. However, the strength of negative associations was larger in magnitude and significantly nonzero across networks (**Figure 2D**), with the exception of the Limbic Network. These results align with previous findings indicating strong decreases in whole-brain functional connectivity as progesterone concentrations increase across the cycle (Pritschet et al., 2019).

**Figure 2.**
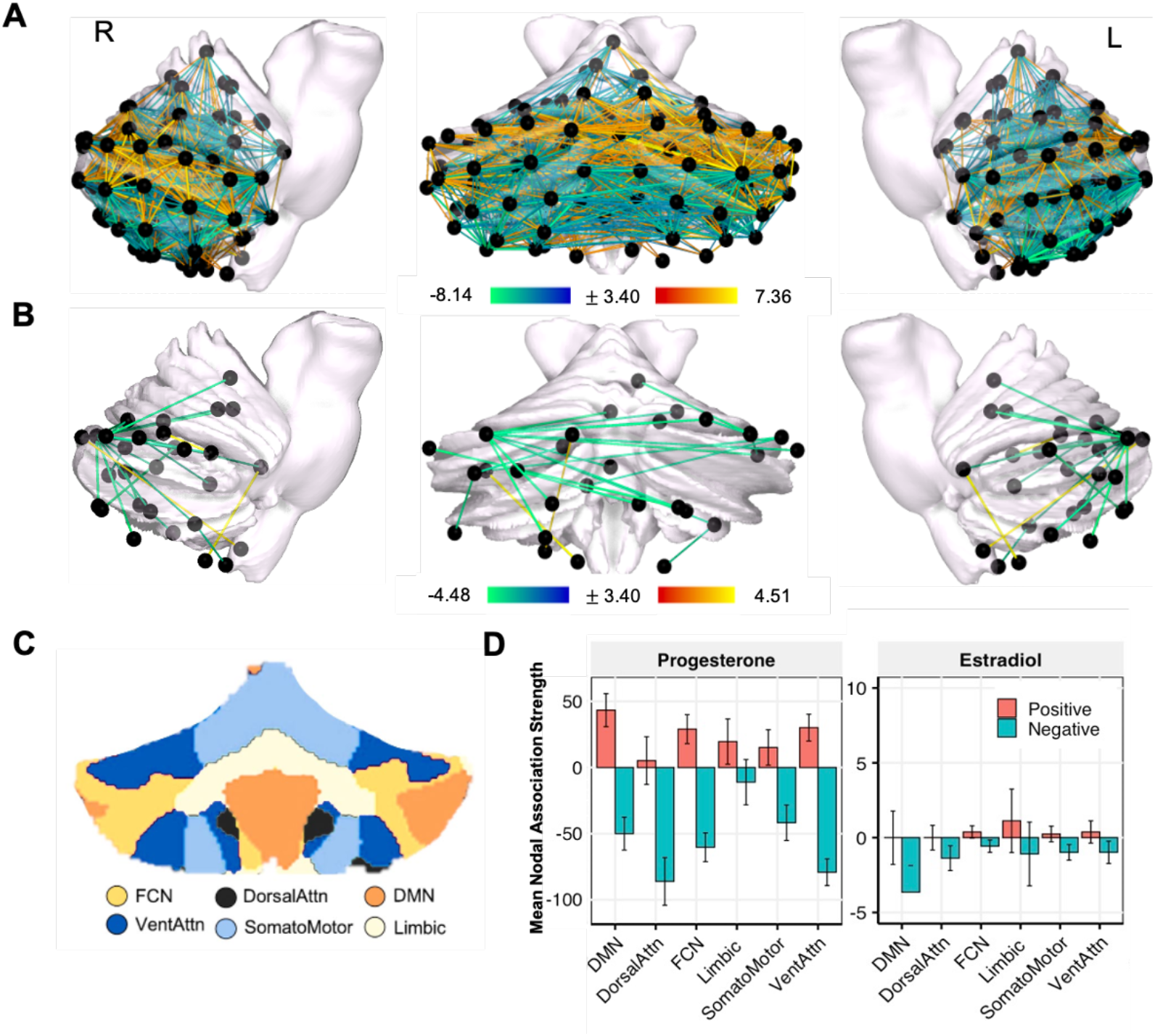
Whole cerebellum functional connectivity at rest is associated with intrinsic fluctuations in estradiol and progesterone. **(A)** Day-by-day associations between progesterone and coherence. Hotter colors indicate increased coherence with higher concentrations of progesterone; cool colors indicate the reverse. Results are empirically-thresholded via 10,000 iterations of nonparametric permutation testing (*p* < .001). Nodes without significant edges are omitted for clarity. **(B)** Day-by-day associations between estradiol and coherence. **(C)** Cerebellar parcellations were defined by Buckner et al. seven-network atlas (2011). Note that the Visual Network is not represented in the cerebellum. **(D)** Mean nodal association strengths by network and hormone. Error bars give 95% confidence intervals. ‘Positive’ refers to the average magnitude of positive associations (e.g. stronger coherence with higher estradiol). Note progesterone had greater associations with edgewise connectivity as reflected in the y-axis range; Abbreviations: DMN, Default Mode Network; DorsalAttn, Dorsal Attention Network; FCN, Frontal Control Network; VentAttn, Ventral Attention Network.

In contrast to our predictions, we observed predominantly negative relationships between estradiol and cerebellar coherence (**Figure 2B**). All cerebellar networks exhibited some degree of significantly negative associations with estradiol (95% CIs not intersecting zero), particularly the DMN and DAN (**Figure 2D**). The Limbic Network was unique again in that it demonstrated a heterogenous response with positive and negative association strengths (**Figure 2D**). These findings suggest that, within the cerebellum, increases in estradiol are predominantly associated with decreases in connectivity, a pattern that differs from the strong positive associations observed across the cerebrum (Pritschet et al., 2019). In general, the negative association strengths related to progesterone were far more pronounced than those observed with estradiol.

### Temporal dependencies between sex hormones and network topology

Given the widespread associations between whole-cerebellar coherence and sex hormones, we examined *topological states* of cerebellar networks to capture the extent of brain-hormone interactions at the network level. Topological states were quantified using common graph theory metrics, including estimates of between-network integration (*participation*) and within-network integration (*global efficiency*; **Figure 3A**).

**Figure 3.**
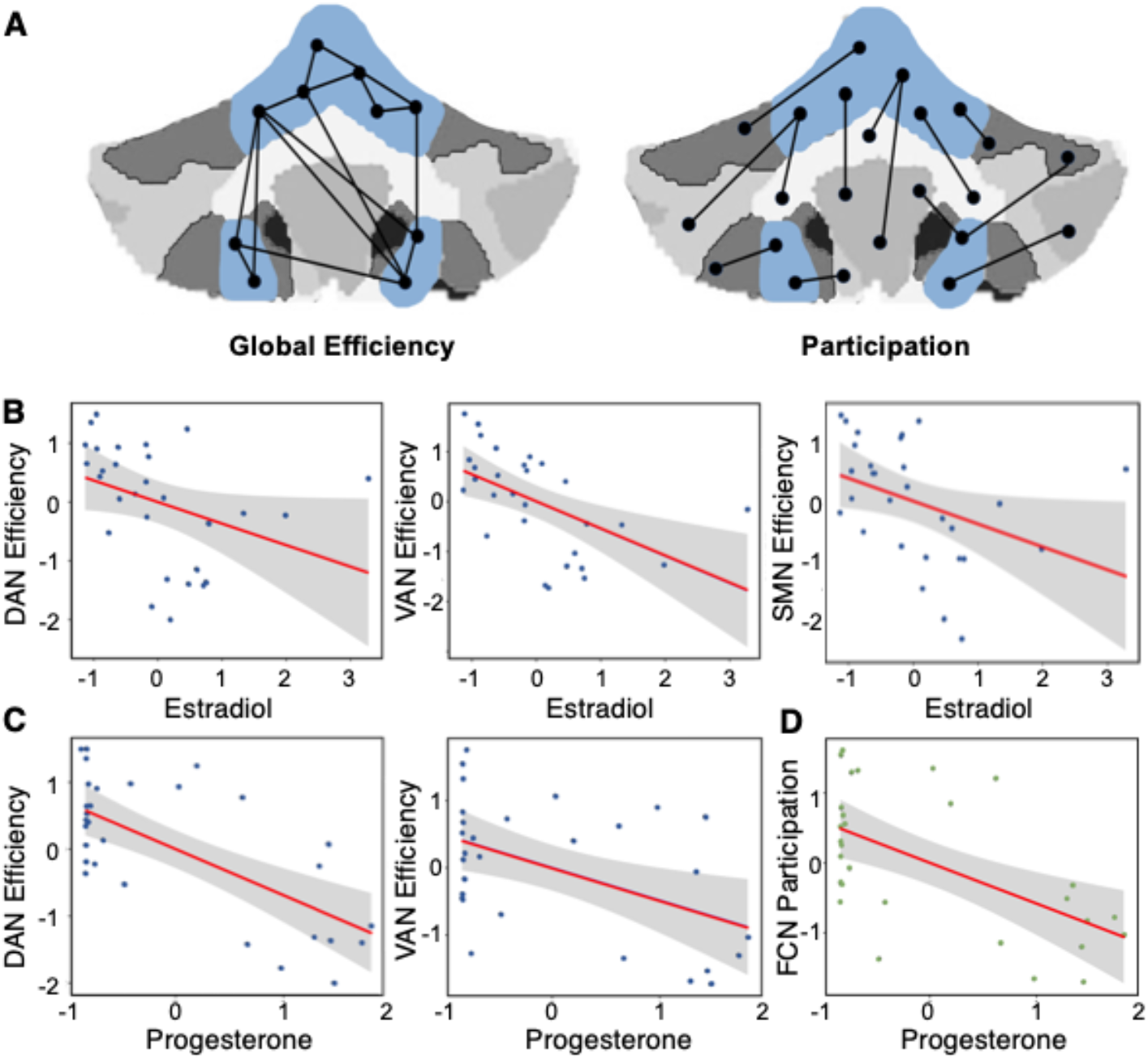
Graph theory metrics reveal relationships between sex hormones and intra- and inter-network integration. **(A)** Illustration depicts common graph theory metrics. Participation coefficient represents a measure of between-network integration, defined as the average extent to which network clusters are communicating with other networks over time. Global efficiency, a measure of within-network integration, reflects the ostensible ease of information transfer across clusters inside a given network. Graph theory metrics for the Somato-Motor Network are shown in blue as an example. **(B)** Scatter plots depict the association between estradiol and DAN, SMN and VAN efficiency (*p* < .05). **(C)** Scatter plots depict the association between progesterone and DAN and VAN efficiency (*p* < .05) and **(D**) FCN participation (*p* < .05). Abbreviations: DAN, Dorsal Attention Network; SMN, Somato-Motor; VAN, Ventral Attention Network; FCN, Frontal Control Network.

To investigate day-by-day relationships between topological states of each network and hormone fluctuations across the menstrual cycle, a series of linear regression analyses were conducted. After initially fitting linear models to the entire dataset, an inspection of residual densities revealed experiment day one as a frequently-poor fit (median absolute deviation > 3); it was therefore removed from the analyses shown here. Remaining data were *Z*-scored and network metrics were residualized on motion (mean FWD) prior to model estimation (*p*-values reported are FDR-corrected at the level of *q* < .05). Regression models revealed that estradiol was associated with global efficiency within DAN (*β* = −.37, *SE* = .16, *t* = −2.36, *p* = .028), VAN (*β* = −.54, *SE* = .16, *t* = −3.51, *p* = .008) and SMN (*β* = −.39, *SE* = .17, *t* = −2.33, *p* = .028; **Figure 3B**). This suggests that the within-network integration (as measured by global efficiency) of major functional brain networks is negatively associated with estradiol across the cycle. Overall model fits were significant for the DAN (F(1, 28) = 5.55, *p* = .028, 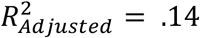, VAN (F(1, 28) = 12.33, *p* = .008, 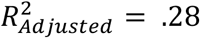) and SMN (F(1, 28) = 5.43, *p* = .028, 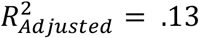). Model fits for the remaining networks were poor and did not demonstrate significant associations with estradiol. These data are in agreement with our edgewise regression analysis depicting decreased whole-cerebellar coherence with increasing estradiol.

Progesterone was associated with DAN (*β* = −.48, *SE* = .15, *t* = −3.29, *p* = .008) and VAN efficiency (*β* = −.39, *SE* = .17, *t* = −2.32, *p* = .028; **Figure 3C**), and both model fits were significant (DAN: F(1, 28) = 10.82, *p* = .008, 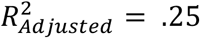; VAN: F(1, 28) = 5.38, *p* = .028, 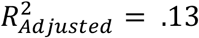). Between-network integration (as measured by participation) for FCN was also associated with progesterone (*β* = −.43, *SE* = .16, *t* = −2.70, *p* = .026; **Figure 3D**), and the model fit was significant (F(1, 28) = 7.28, *p* = .026, 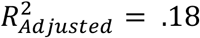). In sum, dynamic changes in progesterone across the menstrual cycle were associated with intra- and inter-network integration of functional brain networks.

## Discussion

In this dense-sampling, deep-phenotyping project, a naturally-cycling female underwent resting-state fMRI and venipuncture for 30 consecutive days, capturing the dynamic endocrine changes that unfold over the course of a complete menstrual cycle. Edgewise regression analyses illustrate robust negative associations between progesterone and cerebellar coherence, and to a lesser degree, negative associations between estradiol and cerebellar coherence. Next, graph theory metrics were used to examine cerebellar network topologies, indicating negative associations between estradiol and global efficiency within the DAN, VAN, and SMN, between progesterone and global efficiency within the DAN and VAN, and between progesterone and participation of the FCN. Together, these results reveal that estradiol and progesterone are associated with cerebellar functional connectivity and network topology, providing insight into the relationship between sex hormones and the intrinsic dynamics of the human cerebellum.

Sex steroid hormones influence *cortical* functional connectivity and network topography, as demonstrated by parallel analysis across the cerebrum (Pritschet et al., 2019). While our predictions for the impact of gonadal hormones on cerebellar network dynamics were analogous to the effect of sex hormones on cerebral networks, our data suggests that day-by-day associations of hormones and cerebellar functional connectivity at rest diverge from that of the cerebrum. While higher concentrations of estradiol were associated with increased connectivity across all major cortical networks, particularly the DMN and DAN (Pritschet et al., 2019), this effect was absent in the cerebellum where estradiol was associated with *reductions* in connectivity across networks, and cerebellar DMN and DAN entirely lacked positive associations. This implies that estradiol might enhance functional connectivity within the cortex while simultaneously decreasing connectivity within the cerebellum. Progesterone’s effects were similar in the cerebellum and cerebrum, demonstrating robust negative associations with coherence in both cases (Pritschet et al., 2019). Notably, the association strengths were considerably higher in the cerebellum (10-fold minimum), hinting that progesterone may have a greater influence on cerebellar coherence. Critically, the mechanisms driving the unique effects of these hormones in the cerebellum have yet to be characterized.

Our results suggest that estradiol has opposing effects across the cerebellum and cerebrum, with minimal negative associations with *cerebellar* coherence but strong positive associations with *cerebral* coherence (Pritschet et al., 2019). This disparity may be attributable to differences in ER subtype distributions. The cerebellum exhibits robust expression of ERβ and almost no ERα, while the cortex has comparable levels of ERα and β (Price & Handa, 2000; Shughrue et al., 1997). ERα and ERβ have similar binding affinities for estradiol (Kuiper et al., 1998; Harris et al., 2002), but the two receptor subtypes diverge in their physiological roles and interactions (Pettersson et al., 1997, 2000; Mosselman et al., 1996; Paech et al., 1997; Saville et al., 2000). Though speculative, the divergent effects of estradiol seen across the cortex and cerebellum could be attributable to the cerebellum’s unique receptor profile. Additional research is needed to definitively link regional differences in ER receptor expression to variability observed at the mesoscopic level of functional networks.

The human cerebellum shows rich expression of PR (Pang et al., 2013) and the human female cerebellum contains higher progesterone concentrations than cingulate cortex (Bixo et al., 1997). If the cerebellum contains relatively higher levels of progesterone than the cerebral cortex in general, this could account for the heightened effects observed between cerebellar coherence and peripheral levels of progesterone. Progesterone potentiates GABAergic activity in cerebellar Purkinje cells (Wilson, 1996), and GABAergic activation by progesterone counters estradiol-induced increases in neuronal excitability (Murphy & Segal, 2000), providing a potential mechanism for our observation of progesterone-related decreases in functional coherence.

Sex steroid hormone’s modulation of cerebellar functional connectivity has implications for understanding human brain function across the lifespan. The cerebellum is involved in a broad spectrum of cognitive functions including learning, memory and decision making (Timman et al., 2010; Mandolesi et al., 2001; Harrington et al., 2004), and age-related declines in cognitive function may be attributable to neuronal changes in the cerebellum. Cerebellar volume declines progressively with advanced age (Tang et al., 2001; Hoogendam et al., 2012; Jernigan et al., 2001; Walhovd et al., 2011) and these age-associated volumetric changes may precede those found in subcortical structures such as the hippocampus (Woodruff-Pak et al., 2010). Sex differences in age-related declines of cerebellar volume have been observed, where midlife women approaching menopause show reductions in cerebellar lobe and vermis volumes relative to age-matched men (Raz et al., 2001; Cho et al., 1999), hinting at a potential role of sex steroid hormones in cerebellar aging. Taken together, our results establish a relationship between sex hormones and cerebellar functional brain network organization. Future work should establish whether there are notable structural differences across the menstrual cycle and whether these mediate the relationships observed between functional connectivity and hormones across cycle.

Sex hormone action in the cerebellum is also implicated in neurodegenerative diseases. Alzheimer’s Disease (AD) is a progressive neurodegenerative disease that exhibits profound sex-differences, with two thirds of sufferers being women (Hebert et al., 2013). Wegiel and colleagues (1999) identified significant reductions in cerebellar volume as a feature of AD pathology. In the progression to AD, the cerebellum undergoes significant morphological alterations, including extensive loss of Purkinje cells, reductions in dendritic spines, and altered dendritic arborization (Mavroudis et al., 2013, 2019). Notably, sex hormone receptor expression (ER and PR) within the cerebellum is highly localized to Purkinje cells (Smith et al., 1989, Price & Handa, 2000). Future studies should investigate whether sex hormones play a role AD-related cerebellar atrophy. Given sex hormones’ ability to shape cerebellar dynamics in a healthy brain, they might also play a role in age- and disease-related cerebellar degeneration.

Limitations of the current study should be considered when interpreting these findings and outlining future investigations. First, the cerebellum is a challenging structure to probe due to its low signal-to-noise ratio and the fact that it contains the vast majority of neurons in only one-ninth of the volume of the cortex (Marek et al., 2018; Schlerf et al., 2014). These challenges result in the cerebellum requiring twice as much resting-state data to achieve the same level of reliability as the cerebrum (Marek et al., 2018). Here, a daily 10-minute resting-state scan was collected from a single individual for 30 consecutive days, providing a robust longitudinal dataset to examine cerebellar functional connectivity. As the amount of data collection needed to achieve intra/inter-reliability is debated within the field (Noble et al., 2017), future work should explore how robust these results are to varying scanning durations.

Second, while the majority of previous cerebellum work has relied on anatomical parcellations, we chose a function-based parcellation to capture the cerebellum’s functional subdivisions. The parcellation we applied outperformed the standard voxel-based approach and other existing cerebellar atlases across measures of node homogeneity, accuracy of functional connectivity representation, and individual identification. However, the parcellation was more accurate when identifying cerebro-cerebellar functional connectivity relative to cerebellar connectivity (Ren et al., 2019), suggesting room for improvement when assessing *cerebellar* coherence. Additionally, a recent publication by Seitzman et al. (2020) proposes that applying a novel ‘winner-takes-all’ partitioning method within the cerebellum produces functionally constrained nodes at an unmatched degree of validity across multiple data sets and anatomical atlases. Our results are reported with respect to one parcellation (Ren et al., 2019); therefore, future work should consider applying multiple parcellations to individual datasets to determine whether robust validity of cerebellar connectivity can be obtained.

Third, our preprocessing pipeline used a spatial smoothing filter (4 mm Gaussian kernel) in effort to achieve a higher signal-to-noise ratio, but application of the smoothing kernel could partially obscure spatial specificity (Behjat et al., 2014, 2015; Reimold., 2006). Note that we repeated our edgewise regression analyses without a smoothing kernel and results largely paralleled findings reported here (see **Figure S1**).

Fourth, resting state scans are highly sensitive to motion. However, in this study motion was limited to fewer than 130 microns per-day on average and robust nuisance signal regression procedures were implemented to reduce motion bias. We additionally took steps to remove day-by-day motion tendencies from our measures of network topology prior to analysis with hormones. That said, replication studies utilizing additional multiple motion correction strategies, such as removal of physiological noise contaminants, are needed to further strengthen these results (Gratton et al., 2020). Note that day-by-day motion was further quantified using a recent filtering approach aimed to reduce high-frequency contamination from motion estimates (Gratton et al., 2020); this approach confirmed consistently low motion throughout the experiment (**Figure S2)**.

Finally, this study densely sampled a single individual over one menstrual cycle, which hinders the generalizability of these findings to a larger population. Follow-up studies that use sparse-sampling methods to investigate cerebellar dynamics in larger samples of women and men across different hormone states (e.g. menstrual cycle, oral contraceptive use, menopause, andropause) will strengthen our understanding of sex steroid hormones role in cerebellar function.

### Conclusion

Over the past 30 years, cognitive neuroscience has established the cerebellum’s integral role in cognition (Diedrichsen et al., 2019; Buckner et al., 2013), dissolving the notion that it is a purely motor-associated region. A parallel literature suggests that the cerebellum is a prominent target of sex hormones (Hedges et al., 2012; Brinton et al., 2008). Here, we demonstrate that endogenous fluctuations in estrogen and progesterone over the menstrual cycle impact the intrinsic network properties of the cerebellum. Thus, when assessing the innate functionality of the human brain, the hormonal milieu must be considered.

## Methods and Materials

### Participant

A right-handed Caucasian female, aged 23 years, underwent venipuncture and magnetic resonance imaging (MRI) scans for 30 consecutive days. The participant had no history of neuropsychiatric diagnosis, endocrine disorders, or prior head trauma. She had a history of regular menstrual cycles (no missed periods, cycle occurring every 26-28 days) and had not taken hormone-based medication in the 12 months prior to the study. The participant gave written informed consent and the study was approved by the University of California, Santa Barbara Human Subjects Committee and all experiments were performed in accordance with relevant guidelines and regulations.

### Study design

The participant underwent daily testing for 30 consecutive days, with the first test session determined independently of cycle stage for maximal blindness to hormone status. The participant began each test session with a behavioral assessment questionnaire followed by an immersive reality spatial navigation task (neither reported here, see Pritschet et al., 2019). Time-locked collection of serum and whole blood started each day at 10:00 a.m. (± 30 minutes) when the participant gave a blood sample. Endocrine samples were collected, at minimum, after two hours of no food or drink consumption (excluding water). The participant refrained from consuming caffeinated beverages before each test session. The MRI session lasted one hour and consisted of structural and functional MRI sequences.

### Endocrine procedures

A licensed phlebotomist inserted a saline-lock intravenous line into either the dominant or non-dominant hand or forearm daily to evaluate hypothalamic-pituitary-gonadal axis hormones, including serum levels of gonadal hormones (17β-estradiol, progesterone and testosterone) and pituitary gonadotropins (luteinizing hormone (LH) and follicle stimulating hormone (FSH)). One 10cc mL blood sample was collected in a vacutainer SST (BD Diagnostic Systems) each session. The sample clotted at room temperature for 45 minutes until centrifugation (2,000 x *g* for 10 minutes) and were then aliquoted into three 1 mL microtubes. Serum samples were stored at −20°C until assayed. Serum concentrations were determined via liquid chromatography-mass spectrometry (for all steroid hormones) and immunoassay (for all gonadotropins) at the Brigham and Women’s Hospital Research Assay Core. Assay sensitivities, dynamic range, and intra-assay coefficients of variation (respectively) were as follows: estradiol. 1 pg/mL, 1–500 pg/mL, < 5% relative standard deviation (*RSD*); progesterone, 0.05 ng/mL, 0.05–10 ng/mL, 9.33% *RSD*; testosterone, 1.0 ng/dL, < 4% *RSD.* FSH and LH levels were determined via chemiluminescent assay (Beckman Coulter). The assay sensitivity, dynamic range, and the intra-assay coefficient of variation were as follows: FSH, 0.2 mIU/mL, 0.2-200 mIU/mL, 3.1-4.3%; LH, 0.2 mIU/mL, 0.2-250 mIU/mL, 4.3-6.4%.

### MRI acquisition

The participant underwent a daily magnetic resonance imaging scan on a Siemens 3T Prisma scanner equipped with a 64-channel phased-array head coil. High-resolution anatomical scans were collected using a *T1*-weighted magnetization prepared rapid gradient echo (MPRAGE) sequence (TR = 2500 ms, TE = 2.31 ms, TI = 934 ms, flip angle = 7°, 0.8 mm thickness) followed by a gradient echo fieldmap (TR = 758 ms, TE1 = 4.92 ms, TE2 = 7.38 ms, flip angle = 60°). Next, the participant completed a 10-minute resting-state fMRI scan using a 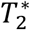-weighted multiband echo-planar imaging (EPI) sequence sensitive to the blood oxygenation level-dependent (BOLD) contrast (TR = 720 ms, TE = 37 ms, flip angle = 56°, multiband factor = 8; 72 oblique slices, voxel size = 2 mm^3^). To minimize motion, the head was secured with a custom fitted foam head case (days 8-30; https://caseforge.co/). Overall motion (mean framewise displacement; FWD) was negligible, with fewer than 130 microns of motion on average each day. Motion was not correlated with estradiol concentrations (Spearman’s *r* = −.06, *p* = .758) but was correlated with progesterone concentrations (Spearman’s *r* = .42, *p* = .020). However, extensive preprocessing steps were taken to minimize motion bias (see **fMRI preprocessing**).

### fMRI preprocessing

Preprocessing was performed using the Statistical Parametric Mapping 12 software (SPM12, Welcome Trust Centre for Neuroimaging, London) in MATLAB. Functional data were realigned and unwarped to correct for head motion and geometric deformations, and the mean motion-corrected image was coregistered to the daily high-resolution anatomical image. All scans were then registered to a subject-specific anatomical template created using Advanced Normalization Tools’ (ANTs) multivariate template construction. A 4 mm full-width half-maximum (FWHM) isotropic Gaussian kernel was applied. Global signal scaling (median = 1,000) was applied to account for transient fluctuations in signal intensity across space and time, and voxelwise time series were linearly detrended. Residual BOLD signal from each voxel was extracted after removing the effects of head motion and five physiological noise components (CSF + white matter signal). Additionally, mean signal from bilateral cerebral cortex within 7 mm of the cerebellum was included as a nuisance regressor to further isolate cerebellar signal (Buckner et al., 2011). Motion was modeled based on the Friston-24 approach, using a Volterra expansion of translational/rotational motion parameters, accounting for autoregressive and nonlinear effects of head motion on the BOLD signal (Friston et al., 1996). All nuisance regressors were detrended to match the BOLD time series.

### Resting-state functional connectivity analysis

Functional network nodes were defined based on the Ren et al. (2019) 100-node cerebellar parcellation and nodes were assigned to Buckner et al. (2011) seven-network cerebellar atlas using a consensus, ‘winner-take-all’ approach. This connectivity-based parcellation was selected because it is superior among existing cerebellar atlases with respect to accuracy of functional connectivity detection, node homogeneity, and individual identification (Ren et al., 2019). Additionally, a 100-node atlas was preferred over other node atlas options (10 and 300 node) because it exhibited more moderate centering of each node and symmetry between the two hemispheres (Ren et al., 2019).

Each day, a summary time course was extracted per node by taking the first eigenvariate across functional volumes (Friston et al., 2006). The regional timeseries was then decomposed into frequency bands using a maximal overlap discrete wavelet transform (Daubechies extremal phase filter, length = 8). Low-frequency fluctuations in wavelets 3-6 (~0.01-0.17 Hz) were selected for subsequent connectivity analysis (Patel & Bullmore, 2016). Finally, we estimated the *spectral* association between regional time series using magnitude-squared coherence: this yielded a 99 × 99 functional association matrix for each day, whose elements indicated the strength of functional connectivity between all pairs of nodes (FDR-threshold at *q* < .05). Note that although the Ren “100-node” parcellation was applied, only 99 nodes were available to be analyzed. We attribute this missing node to the fact that one fell directly on the midline and believe the parcellation combined what were originally two nodes into one.

### Statistical analysis

In order to relate cerebellar-hormone relationships to those observed in cerebral networks, we applied the statistical analyses reported on in Pritschet et al. (2019; see for more detailed methods explanation). In short, day-by-day variation in functional connectivity associated with fluctuations in estradiol and progesterone was assessed through an edgewise regression analysis. Data were *Z*-transformed and edgewise coherence was regressed against the hormonal time series to capture day-by-day variation in connectivity relative to hormonal fluctuations. For each model, we computed robust empirical null distributions of test-statistics (*β*/*SE*) via 10,000 iterations of nonparametric permutation testing and we report only those edges surviving a conservative threshold of *p* < .001 to avoid over-interpretation of effects.

To determine the general direction of hormone-related associations with edgewise coherence, we took the thresholded statistical parametric maps for each model and estimated *nodal association strengths* per graph theory’s treatment of signed, weighted networks. That is, positive and negative association strengths were computed independently for each of the 99 nodes by summing the suprathreshold positive/negative edges linked to them. We then assessed mean association strengths (±95% confidence intervals) in each direction across the various networks in our parcellation.

Here, the 99 nodes were grouped into networks based on their spatial association with large-scale cerebellar functional networks (Ren et al., 2019; Buckner et al., 2011). Through this approach, a total of six cerebral networks are represented in the cerebellum: FCN, DMN, VAN, DAN, SMN, and Limbic Network. The primary Visual Network is not represented in the cerebellum.

Next, we assessed associations between sex hormones and macroscale cerebellar network topologies. Briefly, we computed measures of *between-network* integration (the participation coefficient; i.e. the average extent to which network nodes are communicating with other networks over time) and *within-network* integration (global efficiency; quantifying the ostensible ease of information transfer across nodes inside a given network; **Figure 3A**). To obtain these metrics for each day, the full (99 × 99) FDR-thresholded coherence matrices were subdivided into network matrices as defined by our parcellation. We then computed participation coefficients and global efficiencies for each network using the relevant functions for weighted graphs in the Brain Connectivity Toolbox (Rubinov & Sporns, 2010). Subsequently, a linear regression analysis was conducted. Linear models were initially fit across the complete dataset: an examination of residuals across each network/metric/hormone combination commonly revealed experiment day one as a potential outlier, with a median absolute deviation > 3 relative to the overall residual densities. We therefore removed it and re-ran the analysis: data were *Z*-scored, residualized on motion (mean FWD), and models were re-fit (*p*-values reported are FDR-corrected at a level of *q* < .05).

## End Notes

## Acknowledgements

This work was supported by the Brain and Behavior Research Foundation (EGJ), the California Nanosystems Institute (EGJ), the Hellman Family Fund (EGJ) and the Rutherford B. Fett Fund (STG). Thanks to Mario Mendoza for phlebotomy and MRI assistance. We would also like to thank Caitlin Taylor, Shuying Yu, Evan Layer, and Maggie Hayes for assistance with data collection.

## Author contributions

The overall study was conceived by L.P., S.G. and E.G.J.; M.F., L.P., T.S., and E.G.J. performed the experiments; data analysis was conducted by M.F., L.P. and T.S.; M.F., L.P. and E.G.J. wrote the manuscript; T.S., and S.T.G. edited the manuscript.

## Data/code availability

MRI data and code will be publicly accessible upon publication.

## Supplementary Materials

**Table S1.** Gonadal and pituitary hormones by cycle stage.

|  | **Follicular** | **Ovulatory** | **Luteal** |
| --- | --- | --- | --- |
|  | Mean (SD)<br>*standard range* | Mean (SD)<br>*standard range* | Mean (SD)<br>*standard range* |
| Estradiol (pg/mL) | 37.9 (15.9)<br>*12.5-166.0* | 185.3 (59.0)<br>*85.8-498.0* | 85.4 (26.4)<br>*43.8-210* |
| Progesterone (ng/mL) | 0.2 (0.2)<br>*0.1-0.9* | 0.2 (0.2)<br>*0.1-120* | 9.5 (4.8)<br>*1.8-23.9* |
| LH (mIU/mL) | 5.9 (0.7)<br>*2.4-12.6* | 21.7 (16.4)<br>*14.0-95.6* | 5.5 (2.0)<br>*1.0-11.4* |
| FSH (mIU/mL) | 6.5 (1.2)<br>*3.5-12.5* | 8.1 (3.6)<br>*4.7-21.5* | 4.8 (1.3)<br>*1.7-7.7* |

*Note.* Standard reference ranges based on aggregate data from Labcorp (https://www.labcorp.com/).

**Figure S1.**
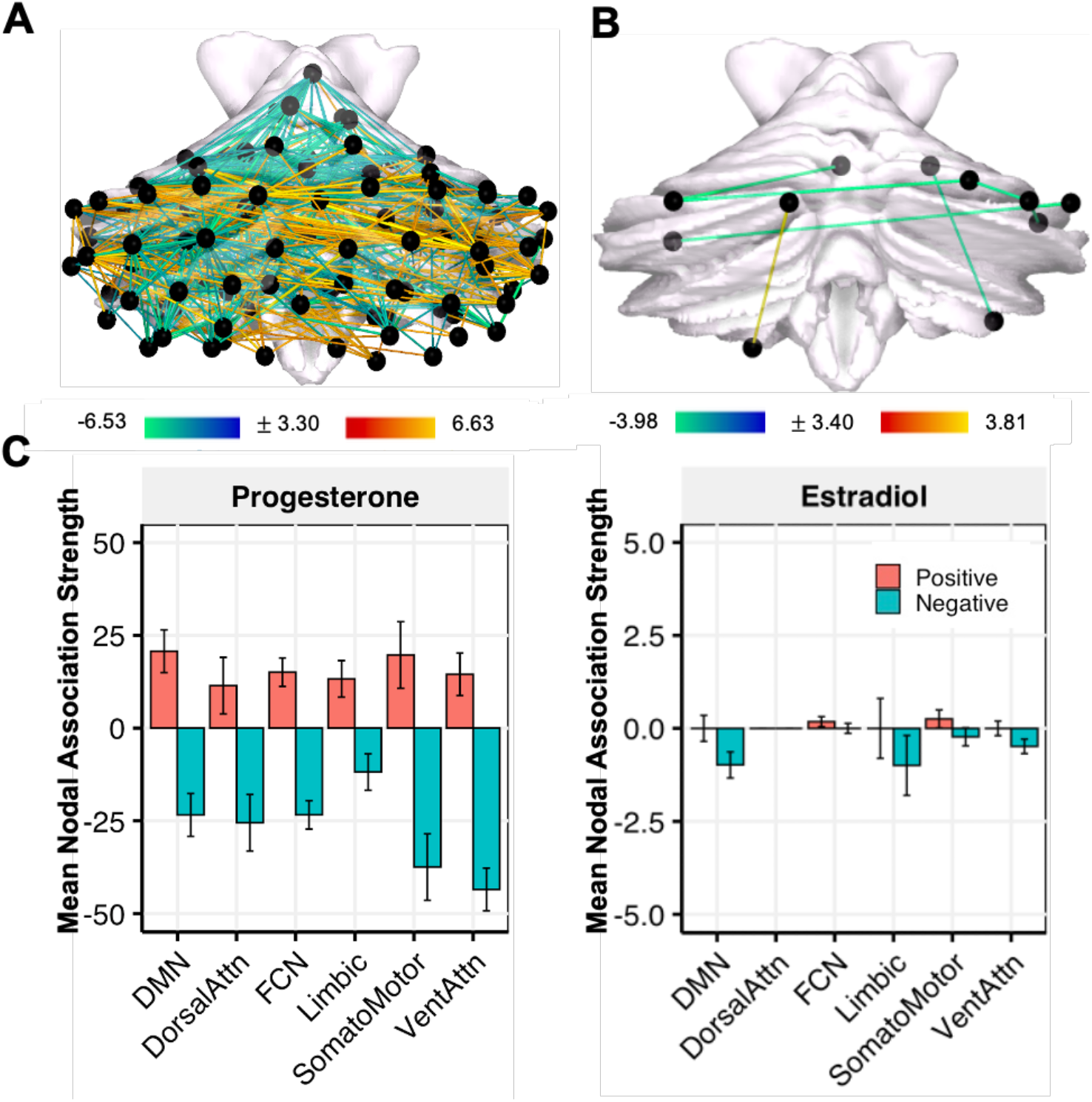
Use of non-smoothed data had minimal impact on brain-hormone associations. **(A**) Day-by-day associations between progesterone and coherence using non-smoothed data. Hotter colors indicate increased coherence with higher concentrations of estradiol; cool colors indicate the reverse. Results are empirically-thresholded via 10,000 iterations of nonparametric permutation testing (*p* < .001). Nodes without significant edges are omitted for clarity. **(B)** Day-by-day associations between estradiol and coherence for non-smoothed data. **(C)** Mean nodal association strengths by network and hormone for non-smoothed data. Error bars give 95% confidence intervals. ‘Positive’ refers to the average magnitude of positive associations (e.g. stronger coherence with higher estradiol). Abbreviations: DMN, Default Mode Network; DorsalAttn, Dorsal Attention Network; FCN, Frontal Control Network; VentAttn, Ventral Attention Network.

**Figure S2.**
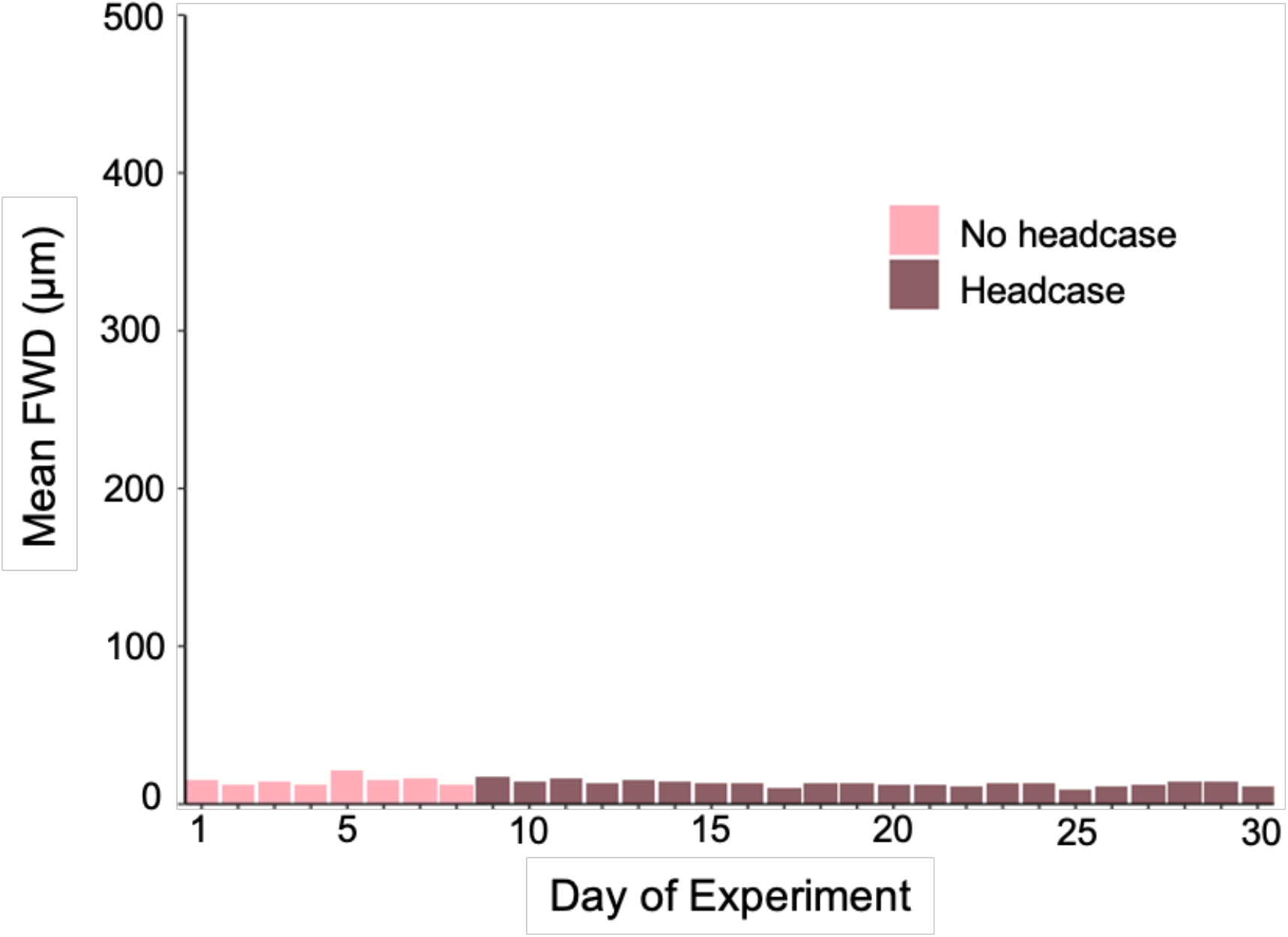
Filtered head motion estimates across the 30-day experiment. Motion was further estimated using a low-pass filtering approach aimed to reduce high-frequency contamination. Mean framewise displacement did not exceed 21 microns, confirming consistently low motion throughout the experiment. Motion on days 1-8 was limited with ample head and neck padding; on days 9-30 motion was limited using a molded headcase custom fit to the participant’s head.

